# A Novel Framework for Quantitative Analysis of Neuronal Primary Cilia in Brain Tissue

**DOI:** 10.1101/2025.05.09.653053

**Authors:** Ali H Rafati, Sofia Rasmusson, Seyedeh Marziyeh Jabbari Shiadeh, Fredrick J Rosario, Thomas Jansson, Carina Mallard, Maryam Ardalan

**Author notes:** Corresponding author: Maryam Ardalan, Department of Physiology, Institute of Neuroscience and Physiology, Sahlgrenska Academy, University of Gothenburg, 40530, Gothenburg, Sweden.

## Abstract

**Background:** Accurate analysis of neuronal primary cilia is essential for understanding developmental processing of neurons. But existing image segmentation methods struggle with staining variability and background noise. To address this, we developed a more robust segmentation and statistical analysis pipeline using an animal model small sample size and with known neuronal microstructure alterations.

**Methods:** Maternal obesity was induced in mice via a high-fat/high-sucrose diet. Hippocampal tissue from 6-month-old offspring of obese and control dams was analyzed. We developed a MATLAB-based pipeline to segment neuronal cilia from z-stack images, applying mathematical transformations and using the Weibull distribution and Bayesian Information Criterion (BIC) to assess group differences

**Results:** The technique segmented cilia despite artifacts, revealing group-specific patterns. Statistical analysis confirmed significant differences, highlighting the method’s robustness over traditional tests, especially with small samples.

**Conclusion:** Our method reliably segments neuronal primary cilia in immune-stained sections with thionin-counter staining and offers a sensitive, assumption-free alternative to traditional statistical tests, ideal for small-sample neurobiological studies

## Introduction

The methods and techniques that are used in automatic image segmentation and mathematical analysis are of special interest in neurobiological studies. Specifically, the application of these techniques in estimation and measurement of micro-structures, such as neuronal primary cilia, are necessary for deep understanding of the underlying mechanism of the neurobiological processes.

Primary cilia are sensory organelles that serve as signalling hubs, detecting and integrating extracellular changes to modulate cellular activity [1]. The neuronal primary cilia have pivotal roles in neuronal proliferation, migration, organization and patterning [2]. Disruptions in neuronal ciliary function during critical developmental periods have been associated with changes in brain morphology and behavioural abnormalities, including cognitive impairment [3, 4]. Primary cilia morphology has shown to rapidly change in response to cellular stressors, including inflammatory [5], hormonal [6] and metabolic [7] signals, which could irreversibly alter primacy cilia function if such changes occur during critical developmental windows [4, 8]. In utero exposure to the abnormal metabolic environment of maternal obesity negatively affect the development of the fetal brain, predisposing the foetus for future neurodevelopmental disease [9-11]. Emerging evidence suggests that maternal obesity is associated with disrupted regulation of neuronal primary cilia in neonatal mice [8]. Investigating these processes using mathematical and computational approaches could provide deeper understanding of the underlying mechanism and function of the primary cilia [12, 13] [14]. Recently, there has been an emerging interest in segmenting and performing 3D reconstruction of neuronal cilia [15] from immuno-stained sections [16]. However, it is not clear whether these methods provide a general solution for image segmentation from brain sections with staining variation and thus, prominent background noise due to staining artefacts. In the next step, implementing so-called mathematical morphology that performs image clustering analysis, the distribution and orientation of neuronal primary cilia from consecutive images could be read and detected more accurately [17]. However, this method does not use mathematical transformation to incorporate pixel identification and filtering, which is problematic because failure to include pattern analysis limits the ability to generate meaningful plots. To overcome this problem, statistical differences can be detected using tests based on the Weibull distribution [18] and by calculating the maximum likelihood estimation in combination with the Bayesian information criterion (BIC) (see also Schwarz criterion) could be applied to compare groups [19, 20].

The current study addresses the challenges of analysing 3D images of microstructures, specifically neuronal primary cilia, in immunostained sections with counter staining. The research focuses on three key aspects:

A. Segmenting neuronal primary cilia from immunostained sections with Nissl counterstaining in z-stack images;
B. Characterizing the spatial patterns of neuronal primary cilia within z-stacks and identifying differences between two groups;
C. Statistically validating these pattern differences using appropriate methods.

We aimed to understand the importance of the classical statistical tests with their assumptions to the small sample size for this type of image analysis and if they are useful in this regard. Accordingly, we developed a method to segment and transform primary cilia pattern data using mathematical equations. By modeling the patterns with a Weibull distribution and applying the Bayesian Information Criterion (BIC), we were able to distinguish between groups with greater sensitivity than traditional methods. This approach reduces reliance on assumptions of normality or homogeneity of variance and is better suited to small-sample image analysis

## Material and Method

### Animals

All experimental procedures were approved by the Institutional Animal Care and Use Committee (Protocol # 00320) of the University of Colorado-Anschutz Medical Campus. A validated mouse model of maternal obesity, mirroring key human features was used [10]. Twelve-week-old proven breeder C57BL/6J females were fed either a control diet (10% fat) or an obesogenic diet (40% fat plus 20% sucrose solution, HF/HS) and all offspring received the control diet. At 6 months, tissues were collected from 12 offspring (6 obese, 6 control; both sexes). One pup of each sex from each litter was used for each offspring outcome studied.

### Tissue Preparation and Immunohistochemistry and Image Capturing

At 6 months of age, mice were perfused with saline followed by 4% paraformaldehyde (PFA). Brains were post-fixed in 30% sucrose for 72 hours, then frozen using isopentane. Sagittal sections (40 µm) were cut using a cryostat (Leica CM 3050 S), with every 8th section selected for analysis using systematic random sampling. Sections underwent immunohistochemistry for Adenylate Cyclase III (ACIII) to label primary cilia. Free-floating sections were pre-treated in target retrieval solution at 85°C, blocked with 3% H_2_O_2_, permeabilized with PBS-T, and incubated overnight at 4°C with rabbit anti-ACIII (1:10,000). After PBS-T washes, sections were treated with biotinylated goat anti-rabbit secondary antibody (1:250) and ABC solution (Vector Laboratories), then visualized with DAB and counterstained with 0.25% thionin. Slides were dehydrated in ethanol, cleared with xylene, and cover slipped. A 100×oil immersion objective lens under light microscope modified for stereology with a digital camera (Leica DFC 295, Germany) and newCAST™ software (Visiopharm, Hørsholm, Denmark) was used to capture images of ACIII-thionin stained.

### MATLAB-Based Image Processing and Statistical Pattern Analysis of Neuronal Primary Cilia in ACIII-thionin-Stained Z-Stack Images

We used three sections per animal in form of z-stack from hippocampal CA1 pyramidal cell layer (CA1.P) to establish the method by using the real data besides simulation and applying the mathematical equations. The implemented codes in MATLAB (R2023b) which includes the neuronal primary cilia image segmentation, image analysis, statistical pattern analysis and simulation are available in supplementary section. Image analysis of the segmented neuronal primary cilia was conducted using the developed equations, which were applied to the coordinates of the segmented cilia. These equations transform the ciliary coordinates into a defined pattern, as illustrated in the simulation section of Figure 2. In the next step, we applied statistical pattern analysis using the Weibull distribution, with model significance evaluated via the Bayesian Information Criterion (Schwarz Criterion) for comparison between the offspring of control and obese dams.

## Results

### Image Processing and Neuronal Primary Cilia Segmentation

By implementing a MATLAB code, the images from the z-stack were transformed into multiple frames (∼15 images in the middle with highest image quality) and then segmented frame by frame by applying the following equations. The obtained images were converted to double precision, with pixel values normalized to the range [0, 1] that is indicated here as (**I**) and then following steps were implemented:

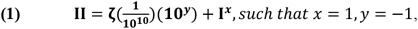

The values of x and y are fixed as given, however if there is image background noise due to tissue staining, these values could be set at the *x* = [0.6 1.4] and *y* = [−0.8 −1]. The details of setting the value are provided in the supplementary section.

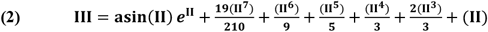

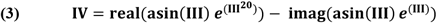

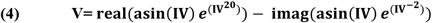

Segmentation of neuronal primary cilia using two parameter sets differing in x-values are shown as an example in **Figure.1**.

**Figure. 1.**
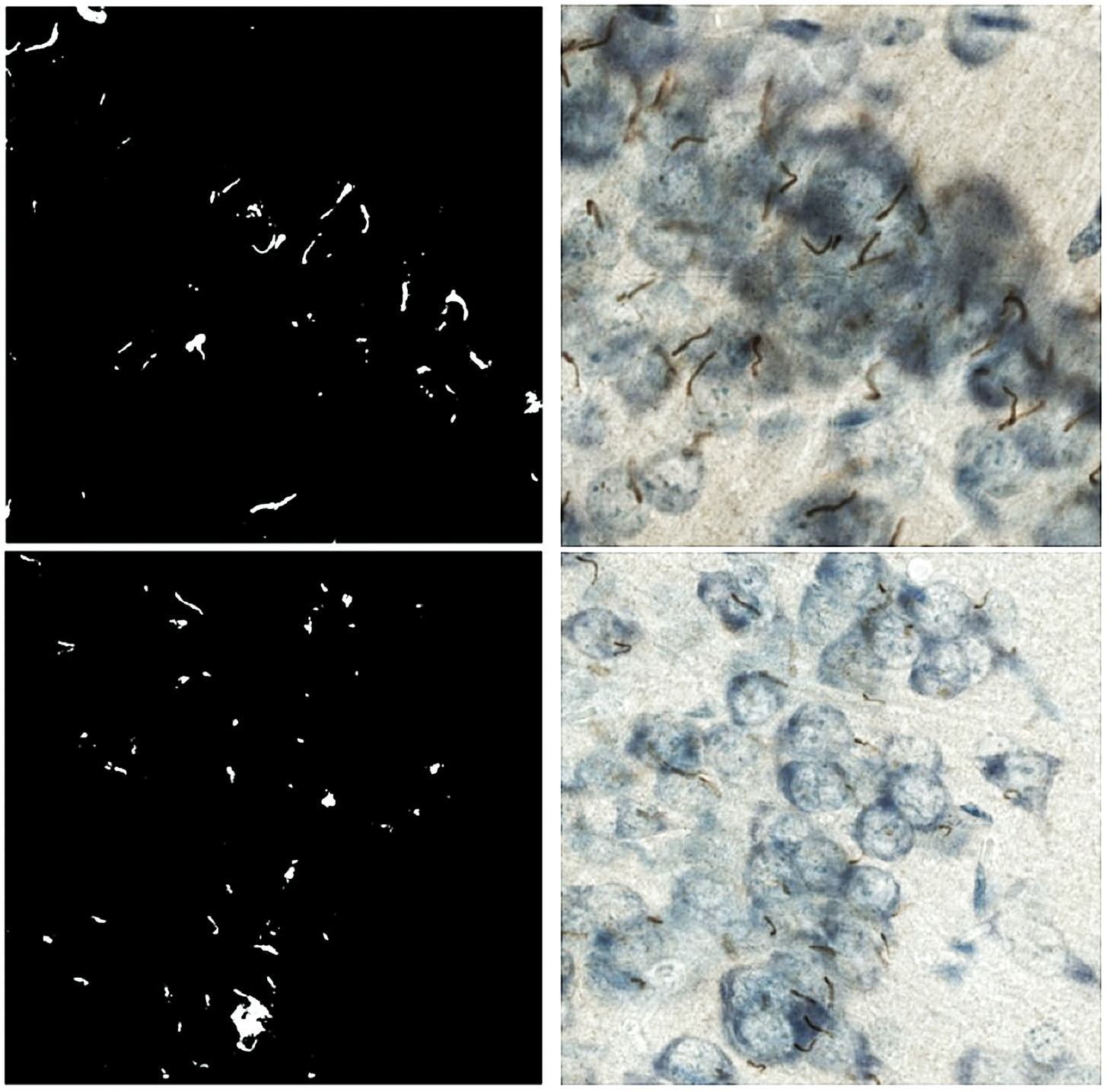
The image processing and neuronal primary cilia segmentation is presented with two examples that differs in x-values. The top, left image is segmented with 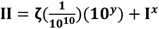, *such that* ***x*** = **0. 8**, *y* = −1 due to straining background noise from the top, right image. While the bottom, left image is segmented with 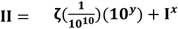, *such that x* = 1, *y* = −1 from the bottom, right image.

### Image Analysis on the Segmented Neuronal Primary Cilia

In this step, we displayed the segmented binary primary cilia. The x, y coordinates of the segmented binary primary cilia in a set ‘*S*’ that should meet these requirements by defining the *S* = {(*x, y*)| *IA*(*x, y*) ≠ ±∞ & *IA*(*x, y*) ≠ 0} that are used to apply the following equations for image analysis:

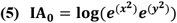

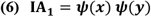

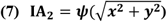

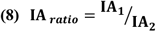

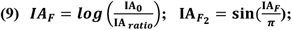

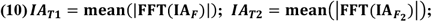

The Fast Fourier Transform(FFT) which is defined as: 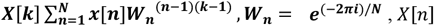 *is input x* = [*x*0, *x*1, *x*2…, *xn*] and *X*[*k*] is output of the ferquncy domain. Next, the simulation of data and real images are shown in **Figure.2** and **Figure.3** respectively.

**Figure. 2.**
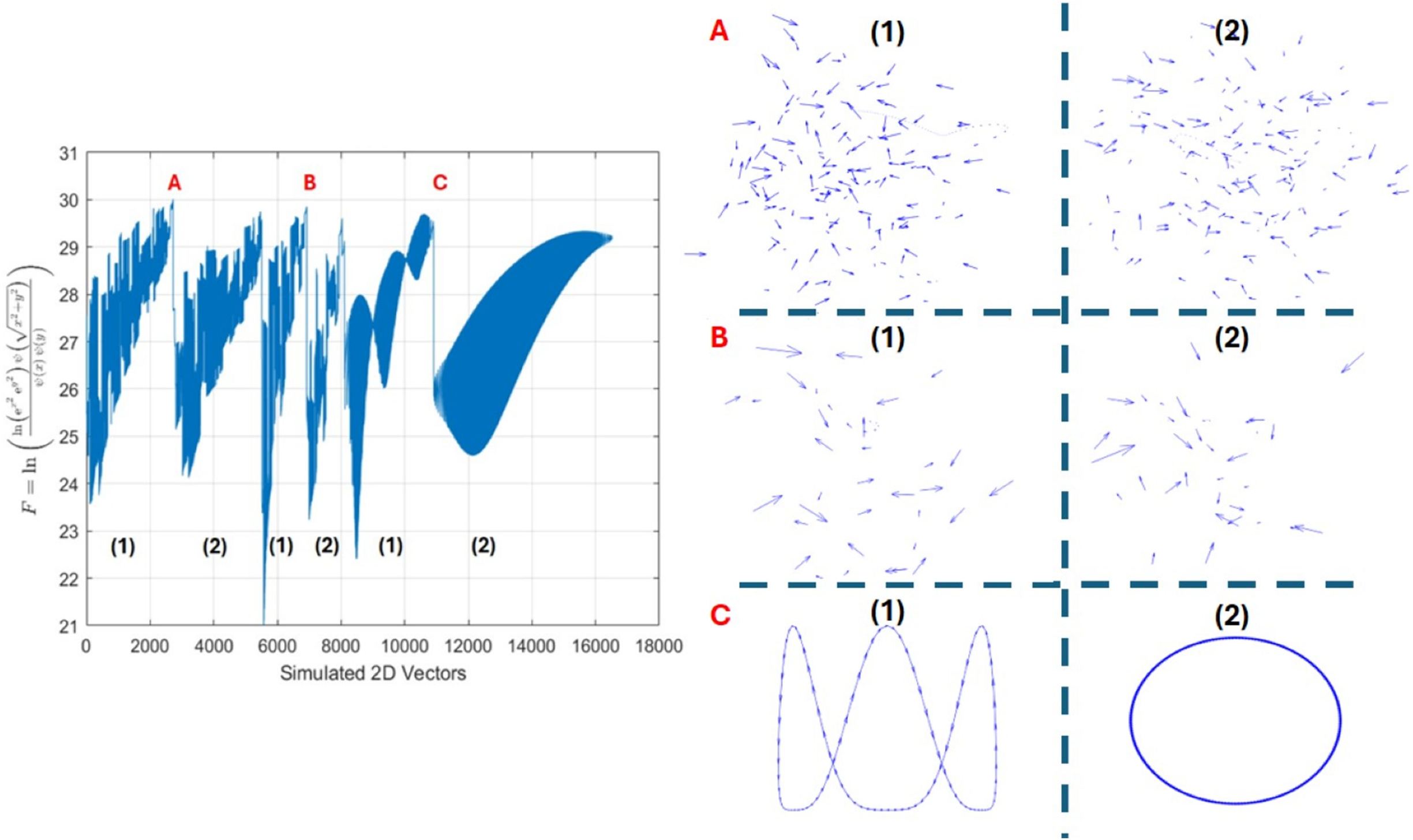
The simulation of the primary cilia distribution was generated on the right side in form of vector-based 2D images (A, B, C) containing two consecutive images for A vs. B. The group C is just a simple different simulation to show how the equations of the ***IA***_***F***_work based on the vector distribution. Then on the left side, the ***IA***_***F***_ (F on the Y-axis) of the images were generated accordingly. It demonstrates how the plot of 2D vectors could look like the cilia distribution. Additionally, the calculated ***IA***_***T*1**_ and ***IA***_***T*2**_ of groups presented as **A**(82,12), **B**(79,9), **C**(102,15) for comparison. These values vary depending on the distribution and density of the vectors (representing simulated cilia). Simulations corresponding to the plots shown on the right are provided in the supplementary.

**Figure. 3.**
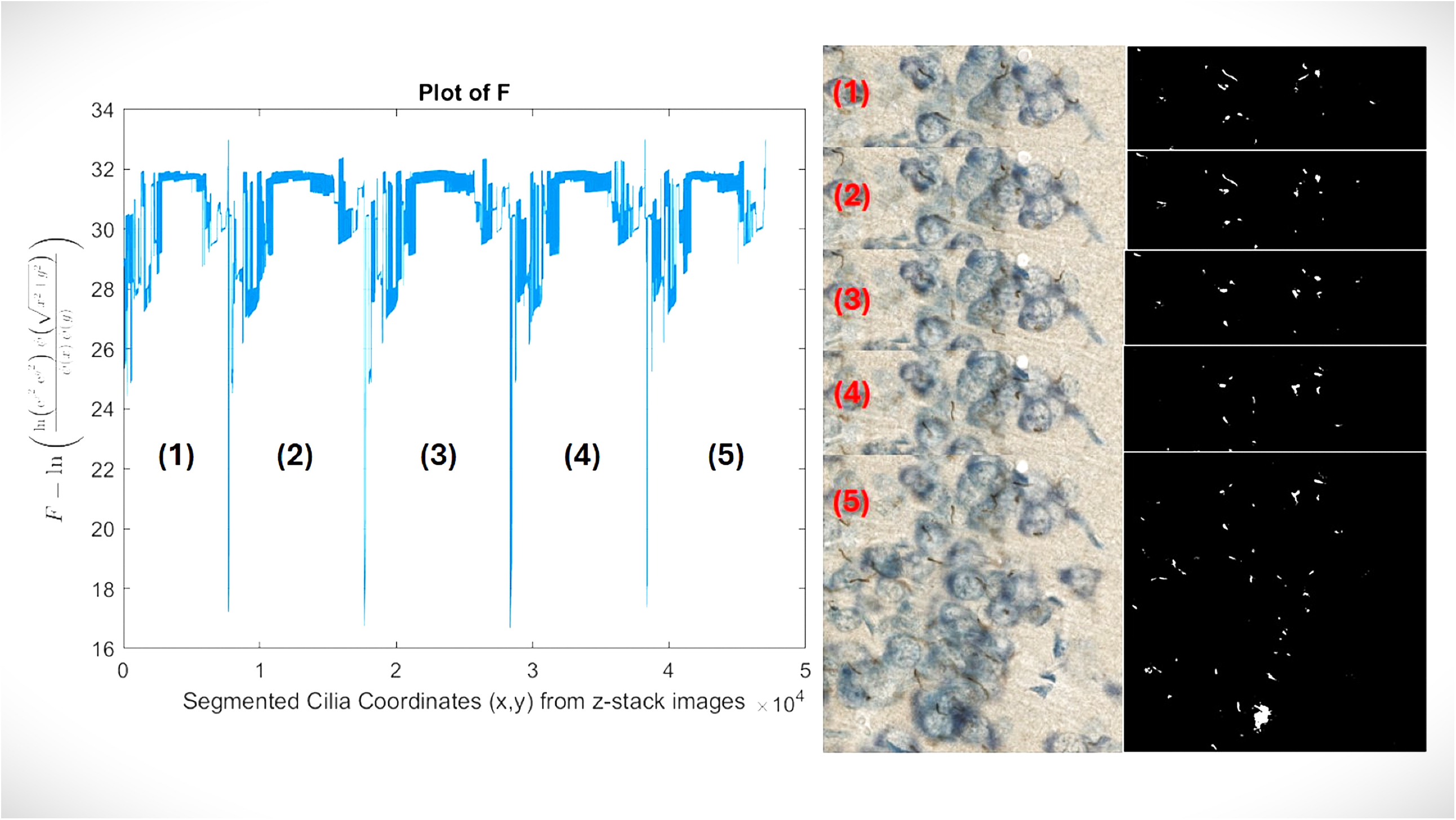
The image analysis on the segmented neuronal primary cilia on five consecutive, original and segmented images that are shown with numbers. The left plot is related to this equation 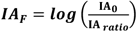 that is shown as **F** on Y-axis, additionally each number (1:5) on the plot is related to the analysis of corresponding segmented image (1:5).

### Description of the Statistical Pattern Analysis of Primary Cilia Applying the Weibull-Distribution and Bayesian Information Criterion (Schwarz Criterion)

In the next step, we performed a statistical pattern analysis of primary cilia using the Weibull distribution, as shown in **Figure 4**. Given that sample sizes in animal studies are typically small and subject to considerable biological variation, traditional statistical tests relying solely on central tendency measures such as the mean or median were not sufficient to meet our analytical needs. These conventional tests are heavily influenced by statistical power and biological variability, which can obscure meaningful patterns in small datasets. Therefore, applying a distribution-based analysis allowed for a more robust assessment of the data. Briefly, we introduced a statistical test that includes the Weibull distribution scale (*λ*) and shape (*κ*) [18], the Weibull negative log-likelihood (*WL*) and its ratio *WL*_*x*/*y*_, Weibull cumulative distribution function (*WCDF*) and its ratio *WCDF*_*x*/*y*_, the data-sorted sum of 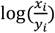 of the given data (X vs. Y group) for comparison in two groups (*S*) or multiple pairwise-groups. Then we obtained the *T*_*x*/*y*_ and *T*_*y*/*x*_ which are the modified and transformed of Weibull negative log-likelihood (*WL*) ratio and are corrected for the ‘n’ that indicates the sample size in each group. The *T*_*x*/*y*_ and *T*_*y*/*x*_ are used to generate the BIC stands for Bayesian information criterion that is also called Schwarz criterion [19, 20]. Additional information on the calculation of the Weibull negative log-likelihood, which can be derived using Equation (11) to estimate the maximum likelihood for comparing two samples, is provided in **Appendix 1** as part of the standard calculation method. Then to test our modified method that includes the Equations (11-16), we applied randomly generated values fitting Weibull distribution of [Scale = 1:3, Shape = 1:3] for sample size = 6:30 over total iteration (∼100) were compared for agreement with the standard method that is described in appendix.1 and available in supplementary. The results are presented in **Figure 5**. The kappa score was used to evaluate the level of agreement between the two tests, with a BIC value greater than 10 considered indicative of significant difference. The MATLAB codes used for conducting the two tests and their comparison are provided in Supplementary.

**Figure. 4.**
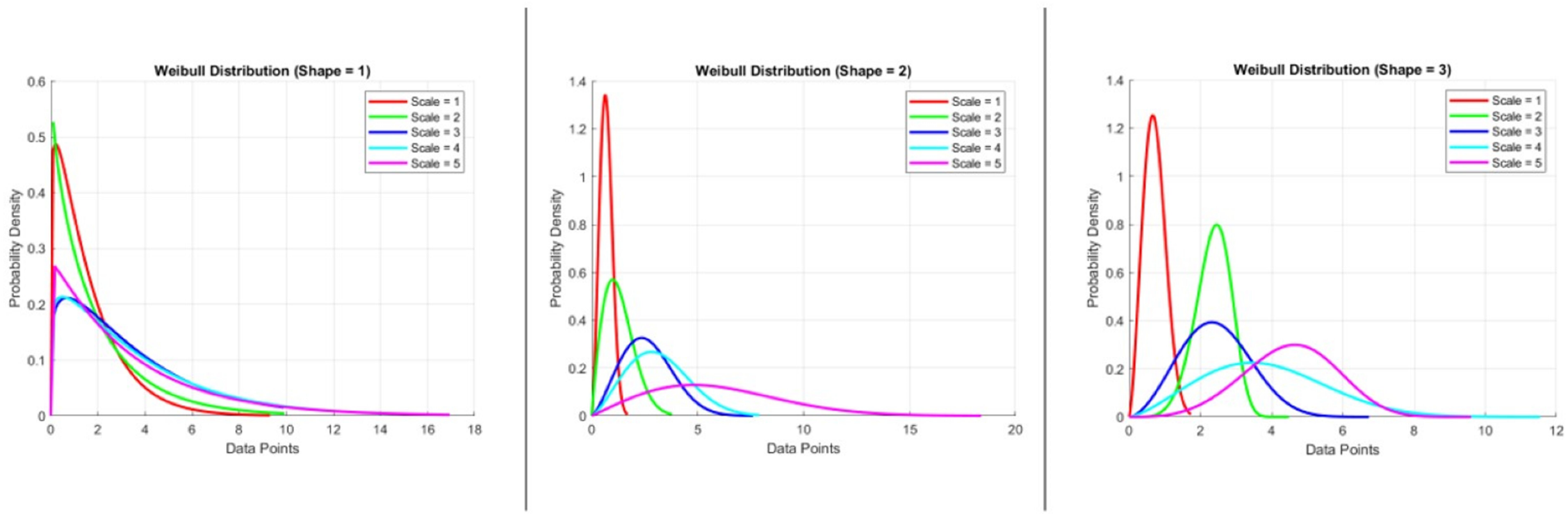
Visualization of Weibull distributions with varying shape parameters (k = 1, 2, 3) and scale parameters (λ = 1–5). Each panel illustrates the probability density function of the Weibull distribution for a fixed shape parameter and a range of scale values. **Left panel:** Shape = 1 (exponential-like distribution); **Middle panel:** Shape = 2; **Right panel:** Shape = 3. As the shape parameter increases, the distribution transitions from a monotonically decreasing curve to a more peaked distribution. The scale parameter shifts the distribution along the x-axis, affecting the spread of data points. These plots demonstrate how the Weibull distribution adapts to different data patterns and can be used to model diverse biological variability.

**Figure. 5.**
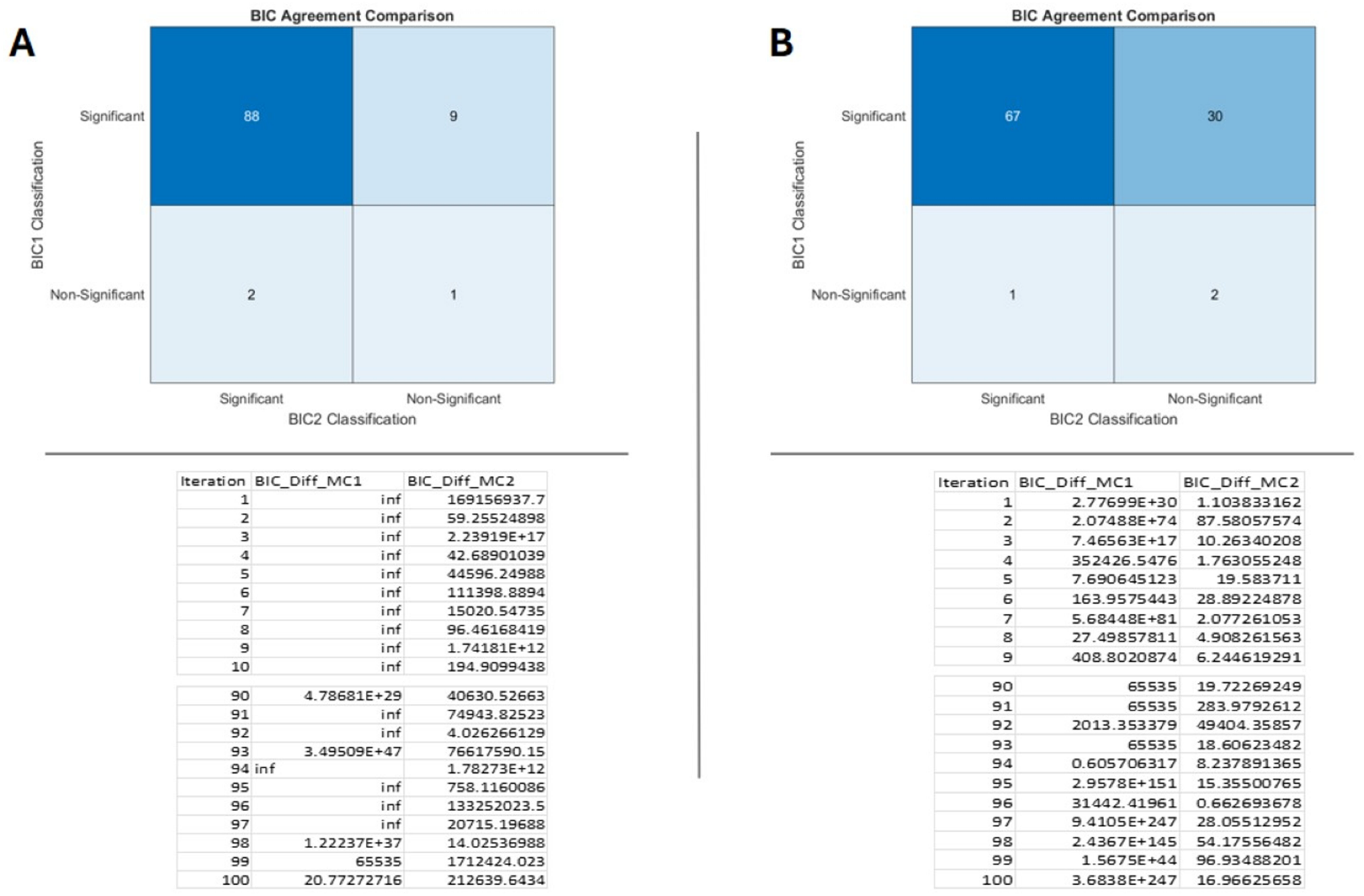
The Weibull distribution of [Scale = 1:3, Shape = 1:3] over total iteration = 100 for sample size n=30 indicted as (**A**) and n=6 indicted as (**B**) that shows the number of significance (BIC >10 was considered significant) in crosstab tables, MC_1 (model calculation_1) represents our described model in Equations (11-16) and MC_2 (model calculation_2) represents the standard BIC calculation using the maximum likelihood ratio from appendix.1. The Kappa= 0.86 for n=30 sample size in (A) and Kappa=0.24 for n=6 sample size in (B). As it shows our test is more sensitive to become significant than the standard model. In the supplementary, the code is available to generate random numbers for 100 iterations for sample size = (6:30) that similar kappa agreement is obtained.

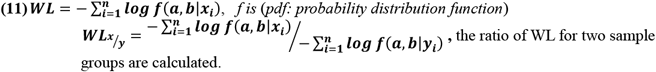

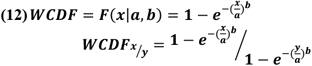

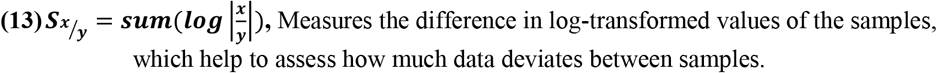

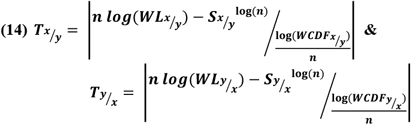

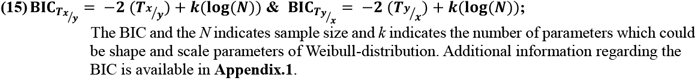

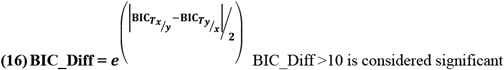

### Intensity-Based Cilia Pattern Characterization and Comparative Statistical Modelling Between Experimental Groups

The analysis on the segmented images yielded the neuronal primary cilia coordinates that the following equation was 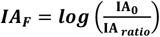 implemented to obtain the plots shown in **Figure.6. A & B** from offspring of control and obese dams. To illustrate the appearance of plots generated from the analysis of fifteen consecutive z-stack images, we performed an one way ANOVA test (not shown in figure 6) and utilized a custom program (available in Supplementary) to automatically and randomly select the plot with the smallest variance difference relative to the rest of the group. However, it required to present the mean of the plots of ***IA***_***F***_ representing the two groups, thus we implemented a step-length sampling of the values in a column that obtained the row-wise mean at each step from all the ***IA***_***F***_ **= [*IA***_***F*1**_, ***IA***_***F*2**_,…***IA***_***Fn***_] contained in certain group, **Figure.6.B**. Next, we demonstrated the statistical analysis of the ***IA***_***F***_ data to compare differences between the two groups. “The statistical analysis was based on the Weibull distribution, applied to the ***IA***_***F***_ **= [*IA***_***F*1**_, ***IA***_***F*2**_,…***IA***_***Fn***_] and structured into two groups ***IA***_***T*1**_ and ***IA***_***T*2**_ as described in Equation (10). The significant difference between the offspring of control and obese dams was demonstrated using the Bayesian Information Criterion (BIC), as detailed in Equations (6–11). A more extensive explanation of the statistical comparison between the two groups is provided in **Figure 7**. Additionally, we performed t-tests and Mann–Whitney tests to evaluate the behaviour of these conventional methods compared to our newly introduced statistical approach. As shown in **Figure 8**, increasing the sample size through replication of the same small dataset could render these traditional tests significant, highlighting their sensitivity to sample size rather than true group differences. In contrast, our proposed method demonstrated greater robustness against sample size variations.

**Figure. 6.**
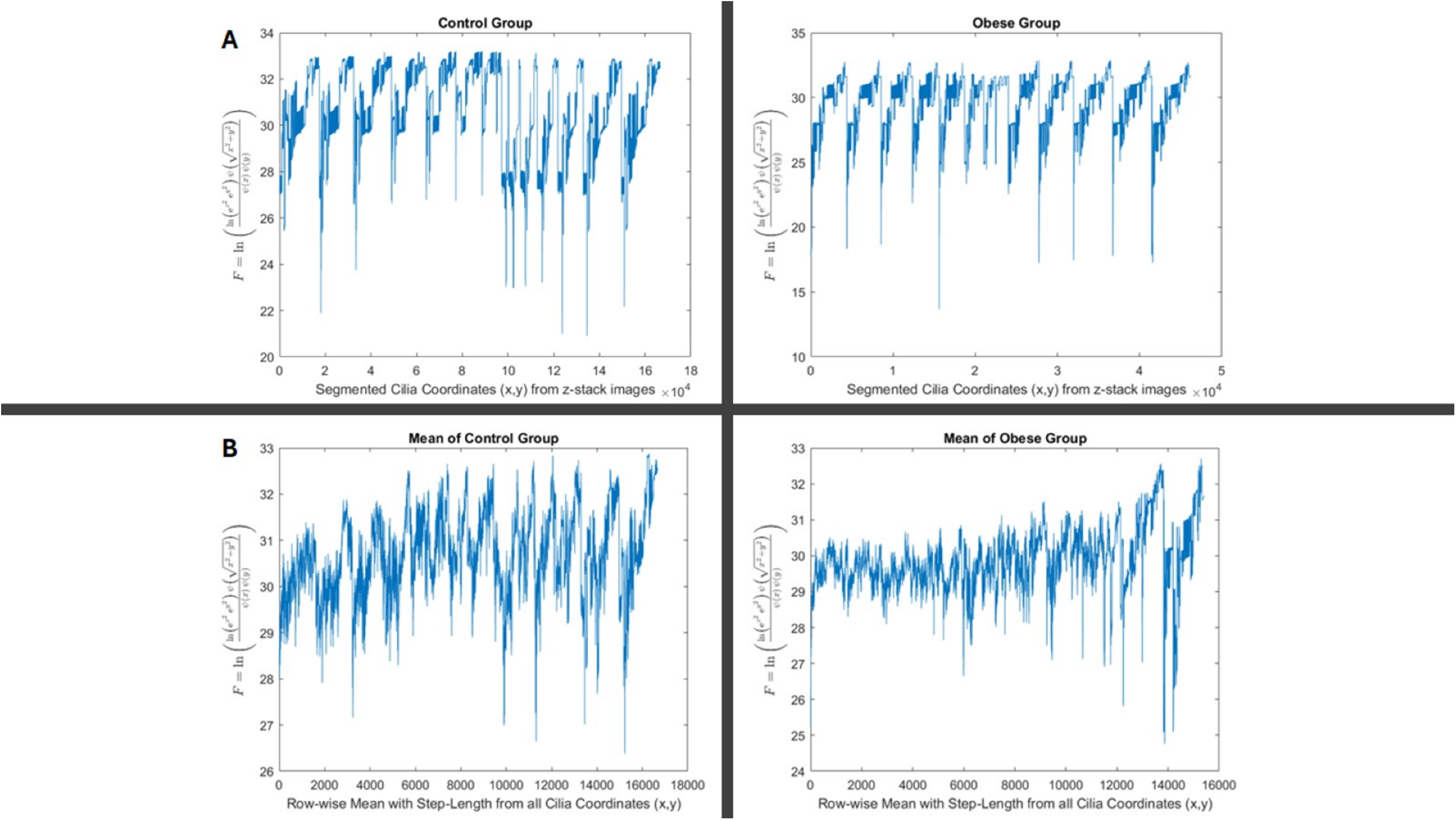
Statistical analysis of segmented neuronal cilia coordinates in control and obese groups using log-transformed intensity analysis. Plots show the calculated 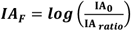 for each individual z-stack image across segmented primary cilia coordinates (x, y) from the control (left) and obese (right) groups. **A**. The images show a random sample chosen from the group with least variance difference, it is example of how consecutive images from a z-stack looks like using the image analysis***IA***_***F***_. Each peak represents variation in the distribution of cilia intensities within the respective image sets. The Control vs. Obese groups show the pattern difference. **B**. Mean ***IA***_***F***_ values were computed using a step-length sampling approach across all cilia coordinates in each group, producing a smoothed, row-wise profile for the control (left) and obese (right) groups. These profiles were used in subsequent statistical comparisons (e.g., BIC and Weibull modelling) to assess group-level differences in ciliary pattern organization.

**Figure. 7.**
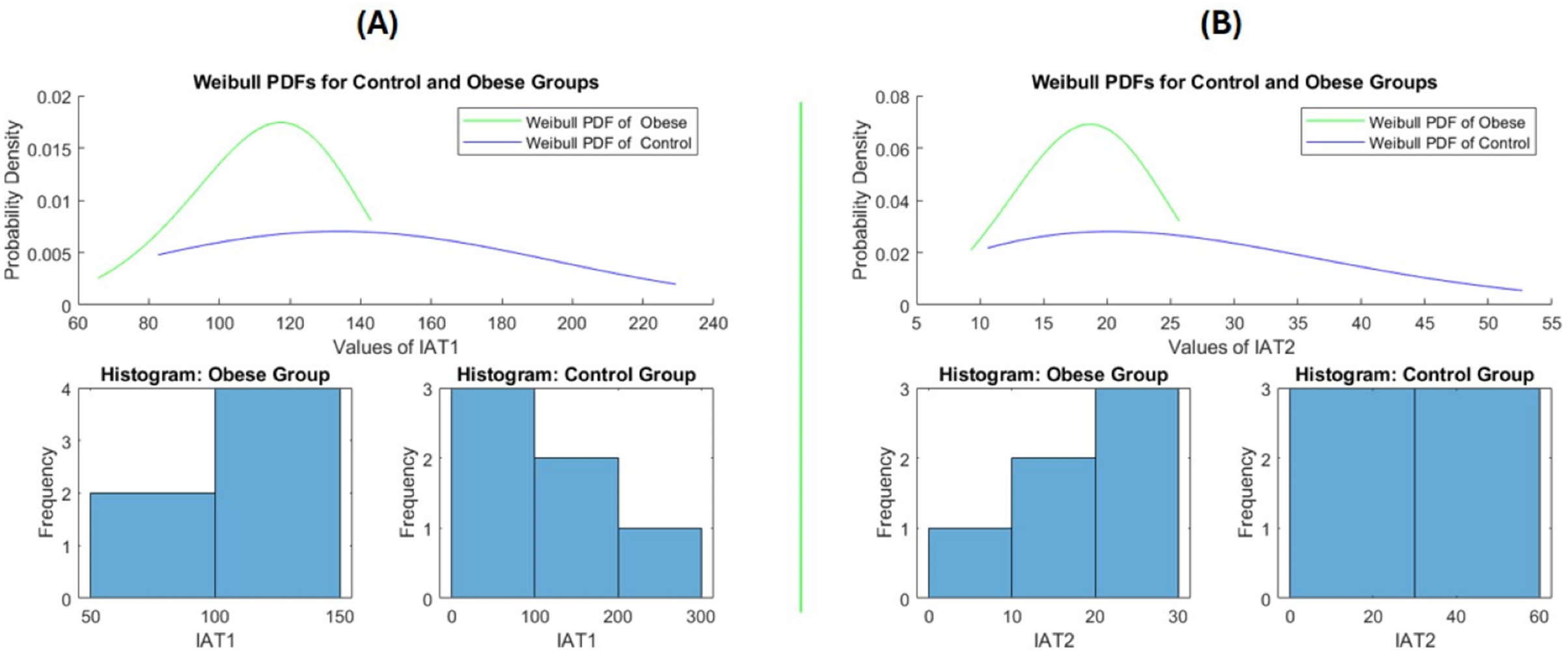
The statistical analysis of Weibull-distribution from the segmented images of neuronal cilia from two groups are shown based on the values obtained from equation (10) ***IA***_***T*1**_ = **mean**(|**FFT**(**IA**_***F***_)|) and 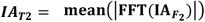.The **(A)** and **(B)** represents the values obtained from ***IA***_***T*1**_ and ***IA***_***T*2**_, respectively. The BIC of both plots showed significant difference between obese vs. control group that was performed by both the introduced statistical test shown in Equations (6:11) and the standard test based on the maximum likelihood estimation shown in in the appendix.1. Thus, the pattern of plots shown in Figure.5 between obese vs. control seems different significantly that is obtained by the mentioned tests.

**Figure. 8.**
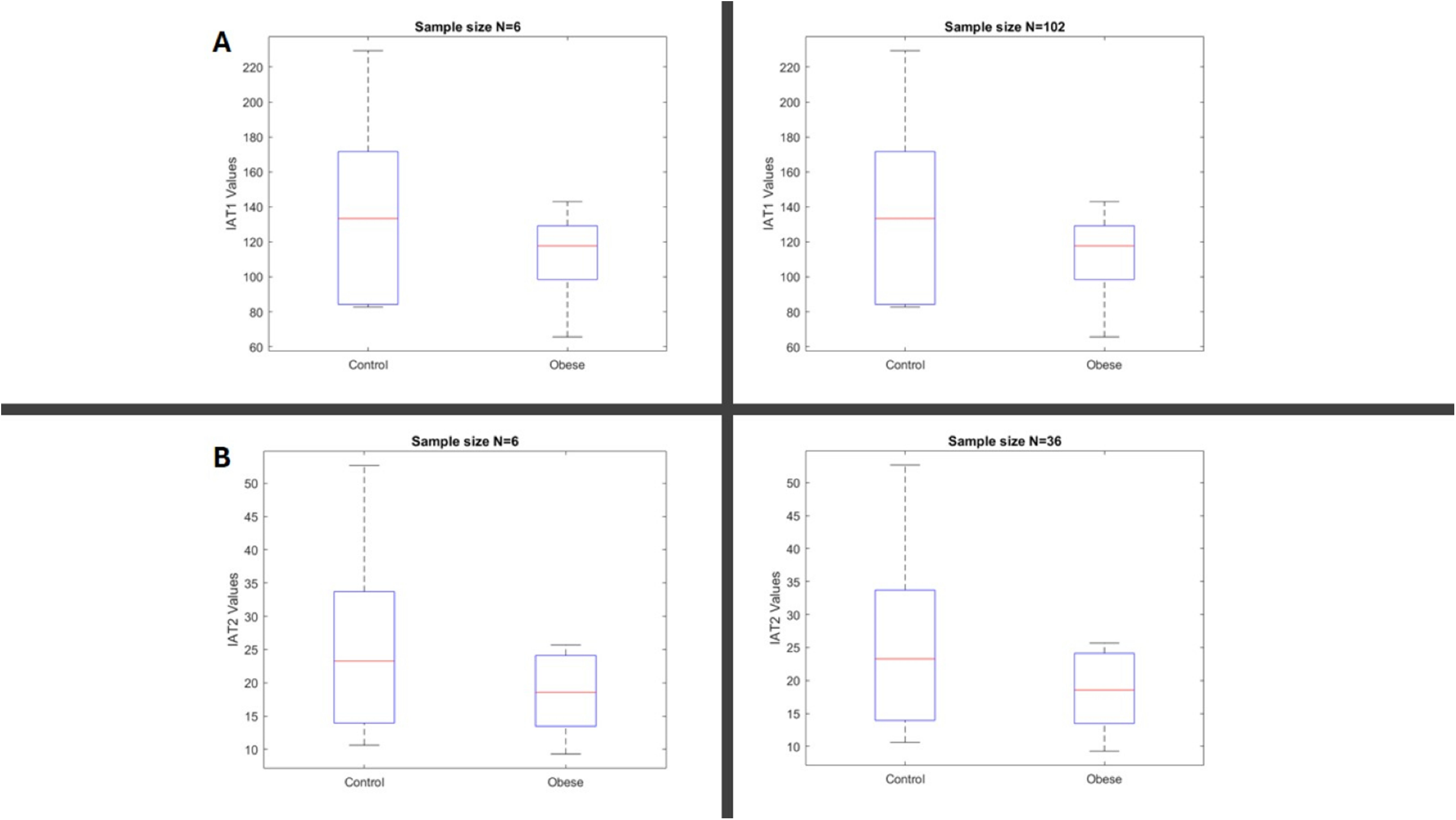
Statistical analysis comparing the obese and control groups was conducted, with the results illustrated in the box plots. Groups A and B were compared based on varying sample sizes. To examine the impact of sample size on the resulting p-values, we replicated identical samples for both groups while keeping the mean and variance constant. When the sample size was small (N = 6), both the t-test and Mann–Whitney test failed to detect a significant difference between the groups. However, when the sample size was increased to N > 30, both tests produced statistically significant results, demonstrating their sensitivity to sample size rather than true group differences. So the **t-test**: for ***IA***_***T*1**_, ***N***=102, ***p***=0.0001and for ***IA***_***T*2**_, ***N***=36, ***p***=0.003, this is while the **Man-Whitney test**: ***IA***_***T*1**_, ***N***=102, ***p***=0.03, and for ***IA***_***T*2**_, ***N***=36, ***p***=0.04.

## Discussion

Primary cilia are critical, hair-like organelles present on nearly all mammalian cells, including neurons and neural progenitors. They play essential roles in neurodevelopment by regulating neuronal migration, axon guidance, and interneuron positioning. Dysfunction of primary cilia has been increasingly implicated in neurodevelopmental disorders, highlighting the importance of accurately studying their structure and distribution in the brain [21].

In this study, we investigated mathematical solutions for segmenting images for analyzing primary cilia in brain sections, immunostained for neuronal ciliary markers, with thionin counterstaining used to provide anatomical reference. By using numerical and mathematical approaches to enable frame by frame segmentation of z-stack images, we developed a mathematical tool that can substantially reduce the image background noise due to staining variability. The mathematical tool in this study is specifically developed for immunostained sections with counterstaining. Compared to other approaches, such as cultured cell models [15] or fluorescent transgenic zebrafish systems [16], immunostained tissue sections sections often present greater challenges due to staining variability and the presence of artifacts. These inconsistencies can introduce significant background noise, making image analysis and signal quantification more difficult and less reliable. In contrast, the image processing method used in this study incorporates parameter-based segmentation, which effectively mitigates the impact of background interference and enhances the accuracy of primary cilia detection, even in complex or noisy tissue environments.

Unlike previous segmentation approaches based on high-resolution, artefact-free images, our method reliably segments primary cilia from conventional brain tissue preparations, thereby expanding the applicability of cilia analysis to a broader range of experimental models. Importantly, by applying a frame-by-frame segmentation across z-stack images, we generated a mathematical framework that captures the distribution and orientation of neuronal primary cilia. The mathematical morphology provided various morphological operators and algorithms that makes the image clustering analysis possible [17]. The resulting two-dimensional coordinate information was transformed mathematically into a vector-based distribution, allowing frequency analysis of repetitive spatial structures. This transformation not only reduces background noise but also emphasizes biologically meaningful patterns of primary cilia orientation and clustering, offering novel insights into the structural organization of neuronal networks. Such analysis is particularly relevant given that abnormal cilia distribution or orientation could disrupt local signalling pathways critical for cortical layering, synaptic development, and ultimately, higher cognitive functions [22]. Therefore, our mathematical modelling bridges image-based observations with biological hypotheses concerning how ciliary alterations may contribute to neurodevelopmental abnormalities. To facilitate group comparisons, we developed a plotting method based on the frequency analysis of transformed 2D cilia coordinates (Figures 2, 3, and 6), offering a quantitative tool for assessing differences in cilia distribution between experimental conditions. However, in order to perform robust statistical comparisons, particularly given the small sample sizes typical of animal studies (n = 6–10), traditional methods relying on mean or median central tendencies were insufficient. To overcome this problem, we applied a Weibull distribution-based statistical framework that models a broader range of distributions than the normality assumptions underlying t-tests. Our statistical pipeline involved point-wise data comparison using the sum of log ratios (S), maximum likelihood estimation (MLE), and the Bayesian Information Criterion (BIC) to assess model fit and group differences. As demonstrated in Figure 7, this approach successfully detected significant differences between offspring of control and obese dams based on cilia distributions. In contrast, standard statistical tests such as the t-test and Mann–Whitney test did not detect significant differences at the original sample size, likely due to limited statistical power. Interestingly, when we simulated a larger sample size (N > 30) through data replication, these conventional tests became significant, highlighting their sensitivity to sample size rather than true biological difference. In comparison, our method, based on MLE and BIC evaluation, proved more sensitive and robust even with small datasets.

Considering the essential role of primary cilia in regulating neuronal migration, interneuron positioning, and signalling during brain development, aberrations captured through altered neuronal primary cilia distribution patterns may serve as early biomarkers for neurodevelopmental disorders. Thus, our approach offers a promising tool for future studies aiming to bridge imaging data with functional neurobiological outcomes, particularly in the context of small-scale, high-variability animal models.

## Conclusion

In this study, we present the development and validation of a novel analytical framework for the reliable segmentation and spatial analysis of neuronal primary cilia in brain tissue. Addressing a key limitation in the field, our method is specifically designed to overcome challenges due to the staining variability and background noise commonly encountered in immunostained sections with counterstaining. By integrating advanced image processing techniques, parameterized mathematical modelling, and a distribution-based statistical approach, we offer a robust and sensitive tool for quantifying primary cilia across heterogeneous samples. To rigorously test the reliability and validity of our method, we applied it to two biologically distinct groups of mice offspring born to either lean or obese dams. This comparative design provided a meaningful test case for evaluating the method’s ability to detect subtle yet biologically relevant changes in ciliary architecture. Our statistical framework, which does not rely on assumptions of normality or homogeneity of variance, proved more sensitive and consistent than conventional statistical tests, especially in the context of variable sample sizes typical of animal studies. Although maternal obesity served as the experimental model in this study, the broader aim was the establishment of a versatile and scalable methodology for primary cilia analysis in the brain. This work lays the foundation for future research into the role of primary cilia in neurodevelopment and disease, providing the field with a reliable tool to investigate early structural biomarkers under diverse experimental conditions.

## Author contributions

The conceptualization, methodology, formal analysis, were performed by AHR. and MA and CM. The animal model was developed and maintained by TJ and FR. Data collection was performed by SR and SMJSH. Writing of the original draft Resources, review & editing, and funding acquisition were provided by the following authors: AHR, M.A, SR and C.M. All the authors reviewed the manuscript.

## Appendix (1)

In this section, we describe the standard statistical method of applying the Weibull-distribution as followed:

1. 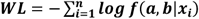, *f is* (*pdf: probability distribution function*)
2. 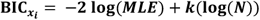, MLE: maximum likelihood estimation is obtained from *WL* based on the shape and scale of the data distribution which is 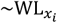. 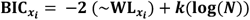, here we omit the *logarithm* as we calculate it in **WL** as *negative logarithmic likelihood*. Similarly, we incorporated the logarithm in equation (14) and not repeated in equation (15). Likewise, the BIC and the *N* indicates sample size and *k* indicates the number of parameters which could be shape and scale parameters of Weibull-distribution.
3. 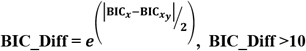 is considered significant, [19].

**Supplementary: There are attached MATLAB codes**.

